# Mitochondrial and insulin gene expression in single cells shape pancreatic beta cells’ population divergence

**DOI:** 10.1101/2020.07.21.213801

**Authors:** H. Medini, T. Cohen, D. Mishmar

**Affiliations:** Department of life Sciences, Ben-Gurion University of the Negev, Beer Sheva 8410501, Israel

**Keywords:** beta cells, gene expression, mitochondrial DNA, mutations, single cells RNA-seq

## Abstract

Mitochondrial gene expression is pivotal to cell metabolism. Nevertheless, it is unknown whether it diverges within a given cell type. Here, we analysed single-cell RNA-seq experiments from ∼4600 human pancreatic alpha and beta cells, as well as ∼900 mouse beta cells. Cluster analysis revealed two distinct human beta cells populations, which diverged by mitochondrial (mtDNA) and nuclear DNA (nDNA)-encoded oxidative phosphorylation (OXPHOS) gene expression in healthy and diabetic individuals, and in newborn but not in adult mice. Insulin gene expression was elevated in beta cells with higher mtDNA gene expression in humans and in young mice. Such human beta cell populations also diverged in mt-RNA mutational repertoire, and in their selective signature, thus implying the existence of two previously overlooked distinct and conserved beta cell populations. While applying our approach to alpha cells, two sub-populations of cells were identified which diverged in mtDNA gene expression, yet these cellular populations did not consistently diverge in nDNA OXPHOS genes expression, nor did they correlate with the expression of glucagon, the hallmark of alpha cells. Thus, pancreatic beta cells within an individual are divided into distinct groups with unique metabolic-mitochondrial signature.

## Introduction

Mitochondrial metabolism is pivotal for the function of all cells, yet it is especially critical for energy demanding tissues, such as brain, muscle and pancreatic beta cells. The hallmark of pancreatic beta cells activity is insulin secretion, which is compromised in type 1 diabetes, and to a lesser extent in type 2 diabetes mellitus (T2DM). Previously it has been shown that insulin secretion was severely impaired upon conditional knockout of mitochondrial transcription factor A (TFAM) in beta cells, thus strongly suggesting that mitochondrial DNA (mtDNA) regulation is essential for insulin secretion (Silva et al. 2000).

It has been previously suggested, that pancreatic beta cells are heterogenous in terms of gene expression, cell surface antigens (Avrahami et al. 2017a), metabolic capacity (Johnston et al. 2016), and rates of insulin synthesis (Avrahami et al. 2017b). Although it has been shown that ATP-stimulated insulin secretion correlate with mitochondrial signalling (Jitrapakdee et al. 2010; Wiederkehr and Wollheim 2012) and relies on active mtDNA regulation (Silva et al. 2000), it is yet unclear whether beta cells are homogenous in regulation of mitochondrial gene expression, and whether such putative variability affects beta cell activity. To address this question, we conducted hypothesis-free analyses of mitochondrial gene expression in three publically available single cell RNA-seq (scRNA-seq) experimental datasets of beta and alpha cells from T2DM and healthy human donors, as well as in three datasets from mouse.

## Results

### Analysis of human scRNA-seq data reveals mtDNA gene expression divergence between alpha and beta pancreatic islet cells

scRNA-seq from human pancreatic beta and alpha cells were analysed in three publicly available datasets (Baron et al. 2016; Xin et al. 2016b; Lawlor et al. 2017) (Fig. 1). The single-cell human transcriptomic Datasets I (InDrops sequencing protocol) contained ∼10,000 human pancreatic cells from four donors isolated from three non-diabetic individuals (ND) and one T2DM patient. Dataset II (Fluidigm C1 sequencing protocol) contained a total of 1492 pancreatic cells from twelve ND and six T2DM donors, and Dataset III (Fluidigm C1 as in Dataset II) contained 638 pancreatic cells from five ND and three T2DM patients. After filtering of Dataset I, while applying quality control measures (taking into account zero inflated reads, a minimum of genes’ number per cell, maximum representation of rRNA transcripts, propensity for doublet cells – see details in Methods), we were left with a total of 2776 cells (1827 alpha cells and 949 beta cells) with ∼2500 informative genes on average per cell. For Datasets II and III, cells with less than 3,000 genes were excluded (due to the different sequencing technology as compared to Dataset I, and the higher sequencing depth of the Fluidigm C1 platform). This resulted in 1396 cells from Dataset II (928 alpha cells and 468 beta cells), and 491 cells from Dataset III (239 alpha cells and 252 beta cells) with ∼5,000 and ∼6,000 informative genes on average per cell for subsequent gene expression analysis, respectively (Table S1). Cells that were called either alpha or beta cells expressed their characteristic transcript, namely either insulin (INS; beta-cells) or glucagon (GCG; alpha-cells) (Dorrell et al. 2011; Baron et al. 2016) (Fig. S1, S2).

**Figure 1:**
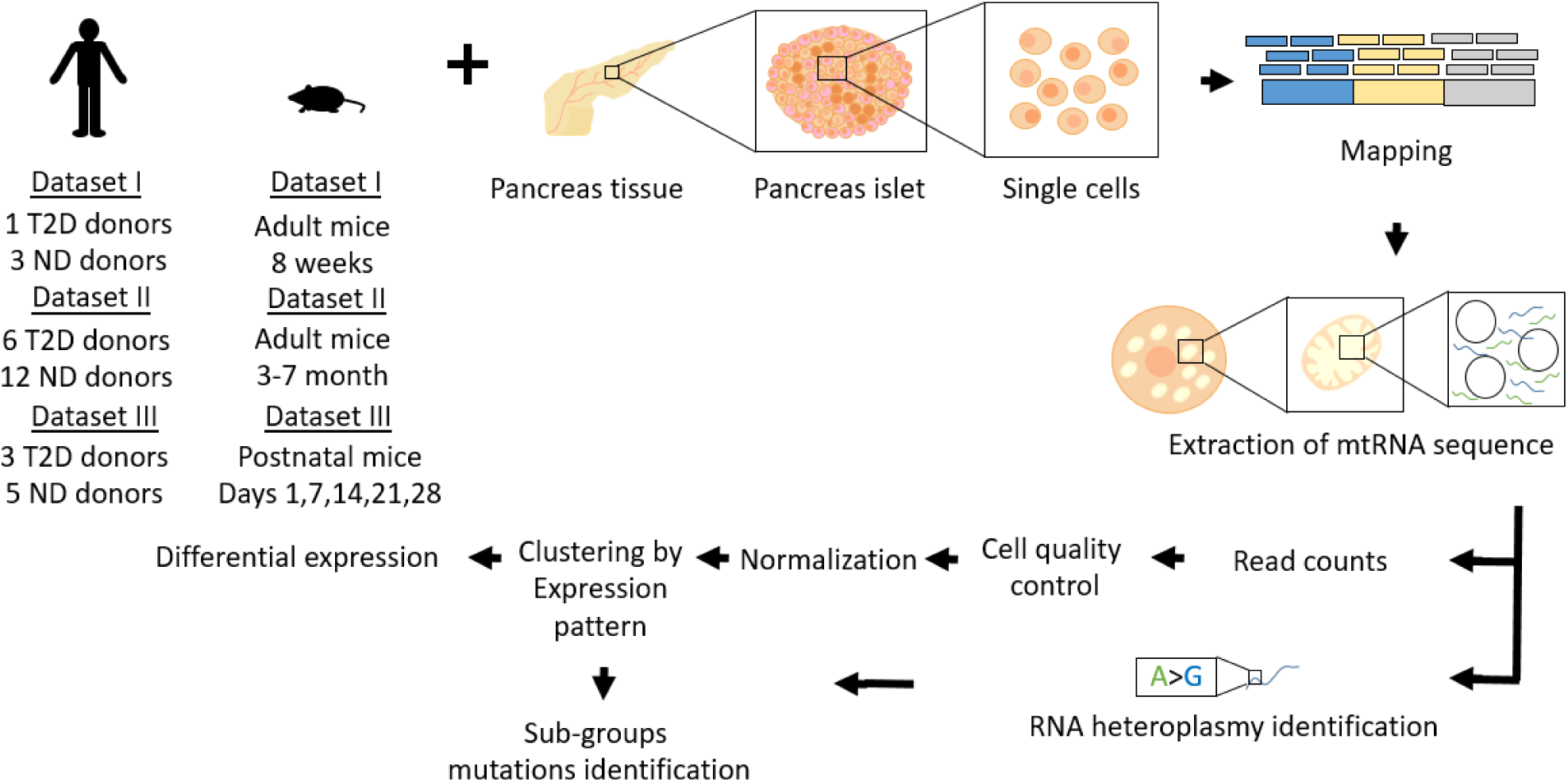
Workflow of scRNA-seq analysis. Hypothesis free scRNA-seq clustering according to mtDNA gene expression. Fastq-files were mapped against the entire genome (GRCh38 for human cells and GRCm38 for mouse cells). After mapping, we calculated read counts, followed by data quality assessment, clustering and differential expression analyses. The RNA mutational heterogeneity data was used to identify mutations that characterize each of the identified cell groups per individual (see Methods).

Comparison of mtDNA gene expression between alpha and beta cells, in healthy and T2DM donors, revealed significantly higher mtDNA transcript levels in beta cells both in healthy and in T2DM patients as compared to alpha cells in all studied datasets (Table S2) (Fig. S3-5). These findings are consistent with known metabolic functional differences between alpha and beta cells in humans (Rorsman et al. 2014). Notably, we could not directly compare healthy and diabetic individuals, as well as assess heterogeneity of the donors in terms of ethnicities and gender, due to the very small sample size (Table S2). The consistency of these findings with previously-published studies encouraged us to continue our analysis further into investigating the subpopulations within each of the tested cell types.

### Human beta cells diverge according to expression of mtDNA and nuclear DNA OXPHOS genes

The function of pancreatic beta cells relies on mitochondrial activity. Nevertheless, it is unclear whether beta cells are homogenous in mitochondrial regulation. As a first step to address this question, we performed hypothesis-free cluster analysis of beta cells from the four donors in Dataset I using Seurat (Butler et al. 2018). This analysis identified two clearly distinct beta cells’ clusters across all tested donors, which differed in mtDNA gene expression per donor (i.e. two groups with either high or low gene expression, designated HE and LE, respectively) (Fig 2A-C). The beta cell subgroups were consistently identified even when analysing all the individuals per dataset grouped together (Fig 2B). Accordingly, the latter analysis revealed that ∼84% of the cells in Dataset I remained in their original subgroups, regardless of the donors, thus further supporting the robustness of these clusters. We next analyzed Datasets II and III to determine the robustness of these sub-groups. To avoid sample size issues, we controlled for sample sizes per donor, and limited our analysis to donors with a minimum of high quality RNA-seq from at least 40 beta cells (Table S1). Despite the small cell number per donor, our findings revealed sharp division into two cell clusters which diverged in the levels of mtDNA gene expression (Fig S6A-B). When analysing all the individuals per dataset together such subgroups were consistently identified as in Dataset I (Fig S6C-D), namely ∼86% and 87% of the cells in Datasets II and III, respectively, remained in their original subgroups, regardless of donors. This supported the robustness of these cell clusters, which were identified regardless of the sequencing platform used. It is worth noting, that this result did not differ between the sequence mapping methods used, i.e. unique or default mapping (Fig S7).

**Figure 2:**
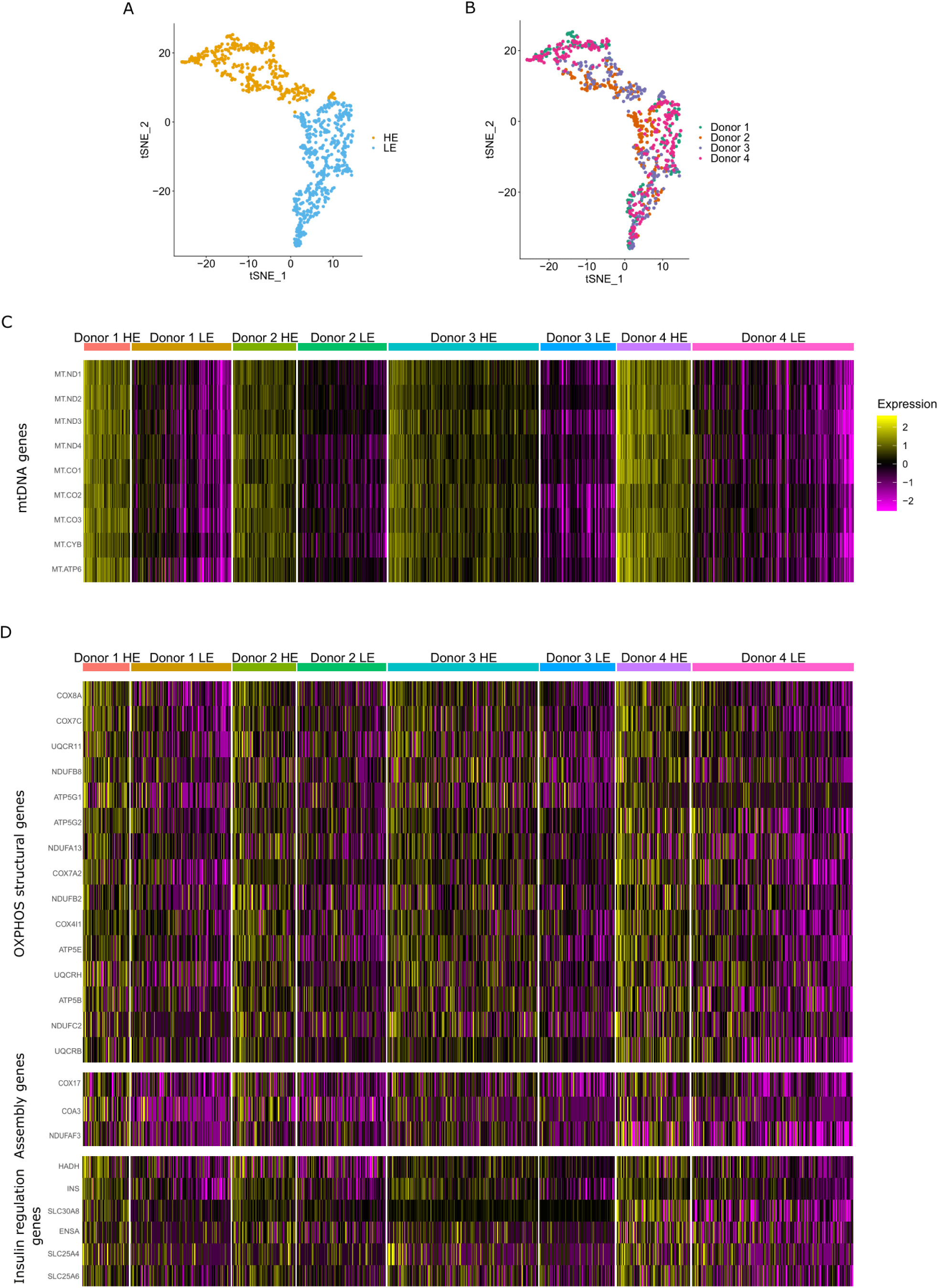
Human mtDNA gene expression analysis revealed two distinct beta cell clusters with either high or low mtDNA gene expression (designated HE and LE, respectively). (A-B) tSNE distribution of beta cells from the four donors (Dataset I) showing two subgroups of beta cells with high and low mtDNA gene expression. Colour codes in (A); subgroups of cells with either high (HE – in yellow) or low (LE – in blue) mtDNA gene expression; (B) Donor identity are colour coded as indicated. (C-D) Heatmap showing the significant differentially expressed genes per cell cluster, per individual (after FDR correction). (C) mtDNA-encoded transcripts, (D) Upper panel -OXPHOS structural genes; Middle panel -OXPHOS assembly genes; lower panel-genes involved in insulin regulation. Colour codes: purple-low expression, yellow-high expression.

Since mitochondrial activities are coordinated between the mtDNA and the nucleus (Barshad et al. 2018), we asked whether the expression of nuclear DNA-encoded (nDNA) genes associates with our observed beta cells populations. As a first step to address this question we analysed genes that consistently differentially expressed between the subgroups across all four donors in a selected set of ∼300 nDNA-encoded proteins which are translated in the cytoplasm and are imported into the mitochondria (Wolf and Mootha 2014; Barshad et al. 2018). This set of genes included all known factors that regulate mtDNA replication, transcription, translation and RNA stability, as well as assembly factors and structural subunits of the mitochondrial OXPHOS system (Pagliarini et al. 2008). Our analysis of Dataset I revealed that the expression of OXPHOS structural genes (complexes I, III, IV and V) consistently diverged between the LE and HE beta cell populations across all four donors (Fig 2B, Table S3). Notably, although to a lesser extent, certain assembly factors of OXPHOS complexes I and III also correlated with these cell populations. To identify differentially expressed genes across individuals in Datasets II and III, we applied the same analysis to the six individuals available from these datasets. The combined analysis of the total samples of beta cells per dataset, revealed expression divergence of OXPHOS structural genes between the HE and LE cellular sub-groups (Fig S8, Table S3), including genes that were consistent between the three tested datasets. Therefore, our results indicate the discovery of novel sub-populations of human pancreatic beta cells that diverge in mito-nuclear OXPHOS gene expression.

### The two beta cell sub-populations diverge in Insulin gene expression

Next, we asked whether our identified beta cell subpopulations associate with the expression of other, additional genes across the human genome. To test for such, we extended our analysis to the entire human transcriptome. Given the set of differentially expressed nDNA genes from the combined analysis (beta cells from all four donors of Dataset I; Table S3) we applied an enrichment analysis to explore which biological processes (GO terms) differentially expressed in the two beta cell subgroups (Table S3). As expected, the analysis revealed that ATP metabolic process and OXPHOS were in the top-ten of the gene list that were upregulated in the HE sub-group of cells (Table S3). We noticed, that the full list of significant processes also included genes involved in insulin regulation and secretion. Strikingly, we found that the cell cluster with higher mtDNA gene expression showed significantly higher expression of INS, encoding the insulin transcript (p<1×10^−50^, Dataset I; FDR correction). To assess whether the differential expression of the insulin regulatory pathway is more prominent than other pathways, we assessed differential expression of selected gene pathways between the HE and LE groups of cells: regulation of insulin secretion, cell proliferation, glycolysis and cell cycle. This analysis revealed, that the HE cells’ group showed significantly high expression of genes involved in regulation of insulin secretion, including the following: Firstly, SLC30A8, encoding a zinc-efflux transporter (zinc transporter 8 (ZnT8)) which mediates uptake of zinc into secretory granules (p<0.05, Dataset I, FDR correction) (Davidson et al. 2014); secondly, SLC25A4 and SLC25A6 which translocate ADP from the cytoplasm into the mitochondrial matrix and ATP from the mitochondrial matrix into the cytoplasm (Gutierrez-Aguilar and Baines 2013); third, ENSA which encodes an alpha-endosulfine, a regulator of the beta-cell K(ATP) channels (p<0.05, Dataset I, FDR correction) (Heron et al. 1999), and HADH gene, a negative regulator of Insulin secretion (Pepin et al. 2010) (Fig 2D). Finally, PTPRN, which participates in the beta cells proliferation pathway and normal accumulation of secretory vesicles (p<1×10^−8^, Dataset I; FDR correction) (Stutzer et al. 2012) was also upregulated in the HE subgroup. Notably, PPP1R15A, an unfolded protein response (UPR) gene, that was previously associated with low INS gene expression levels in mouse beta cells (Lipson et al. 2006; Xin et al. 2018), was upregulated in the beta cells group with lower mtDNA gene expression (LE) (p<0.0032, Dataset I; FDR correction). Notably, the expression of INS was also consistently higher in the HE subgroup of beta cells in Datasets II and III (p<1×10^−5^, Dataset II, FDR correction; p<0.005, Dataset III, FDR correction), thus further attesting for the robustness of this result. It is worth noting, that while examining additional mito-nuclear genes we found that the expression of MEF2D – a transcription factor that was shown to regulate both nDNA and mtDNA gene expression (She et al. 2011), was higher in the HE subgroup (p<1×10^−6^, Datasets II, FDR correction; p<1×10^−11^, Dataset III, FDR correction), thus suggesting an attractive candidate regulator, which explains differences between the HE and LE subgroups. Taken together, our mito-nuclear co-expression analysis strengthen the interpretation that pancreatic beta cells are divided into sub-populations which diverge in mitochondrial gene expression. This divergence is not only limited to mitochondrial activities, but also associates with the expression and regulation of insulin – the hallmark of beta cells’ function. To our knowledge, these results serve as the first demonstration of mitochondrial regulatory involvement in physiologically relevant variability of beta cells activity.

Beta cells heterogeneity was previously mentioned in context of specific antigens and their expression (Dorrell et al. 2016); although the identified subgroups of cells in Dorrell et al are not apparently associated with mitochondrial function, we assessed whether our identified groups of cells correlate with this published sub division of human beta cells. Our analysis did not support correlation between the expression of the antigens identified by Dorrell et al. with our identified sub-groups of beta cells (Fig S9).

### RNA mutational repertoire is elevated along with mtDNA expression

The divergence of beta cells according to the expression of both mtDNA and certain nDNA genes suggests that human pancreatic beta cells are divided into two populations with distinct mitochondrial profiles. Since unlike the nDNA, the mtDNA resides in multiple cellular copies that may differ in sequence (heteroplasmy) we asked whether our observed differential mtDNA gene expression between the two beta cell groups also display differences in mitochondrial RNA (mt-RNA) sequences. Notably, mt-RNA sequence heterogeneity could stem from mtDNA sequence variation, RNA sequence heterogeneity (due to RNA polymerase errors) and RNA modifications, as we recently discovered (Bar-Yaacov et al. 2013; Bar-Yaacov et al. 2016; Safra et al. 2017). To identify mt-RNA mutations with high confidence, we determined RNA heterogenetic mitochondrial mutations with a computational pipeline that utilized individual per-base sequence differences, while employing quality control measures to avoid sequencing errors. As mentioned above, due to low sequence coverage at the non-coding mtDNA region, we focused our analysis on the protein coding mtDNA sequences. Then, we verified that each tested individual had more than a 1000 mtDNA positions with high sequence coverage (>400x). This requirement enabled analysing the sequences generated in Datasets II and III, but not in Dataset I, which displayed lower per base coverage (Datasets I contained on average 100,000 reads for each analysed cell as compared to an average sequencing depth 0.95 ± 0.46 million reads and 34 million reads in Datasets II and III, respectively). While interrogating the repertoire and distribution of the RNA heterogenic mutations we found greater mt-RNA mutational repertoire in the HE group as compared to the LE cell cluster in both Datasets II and III (p<0.005, Dataset II; p<1×10^−16^, Dataset III) (Fig 3A, Table S4). Additionally, we noticed lower number of overlapping mutations as compared to unique mutations of the subgroups (Table S4). Next we divided the mt-RNA mutations into candidate inherited mutations (i.e., shared between alpha and beta cells from the same individual) and ‘others’. As intuitively expected, the percentage of candidate inherited mutations in each beta cells group was found to be higher in the overlapping mutations between the groups as compared to the unique mutations in each group), per individual (Table S4), and the proportion of these mutations was higher in the LE group as compared to the HE group in five out of six individuals. To better understand the functional potential of the mutations in each subgroup, we tested whether RNA heterogenic mutations occurred randomly throughout the mtDNA, per subject. Interestingly, the observed mutational conservation score was lower than expected by chance in both groups, although the LE group had significantly higher score as compared to the HE group in both datasets (p<0.05) (Fig 3B) and a tendency towards higher score in all six tested individuals (Fig S10). Thus, the two beta cells sub-groups differ in mitochondrial mutational repertoire, and in the potential impact of such mutations, suggesting a stronger signature of negative selection acting on mt-RNA mutations in the HE beta cells group.

**Figure 3:**
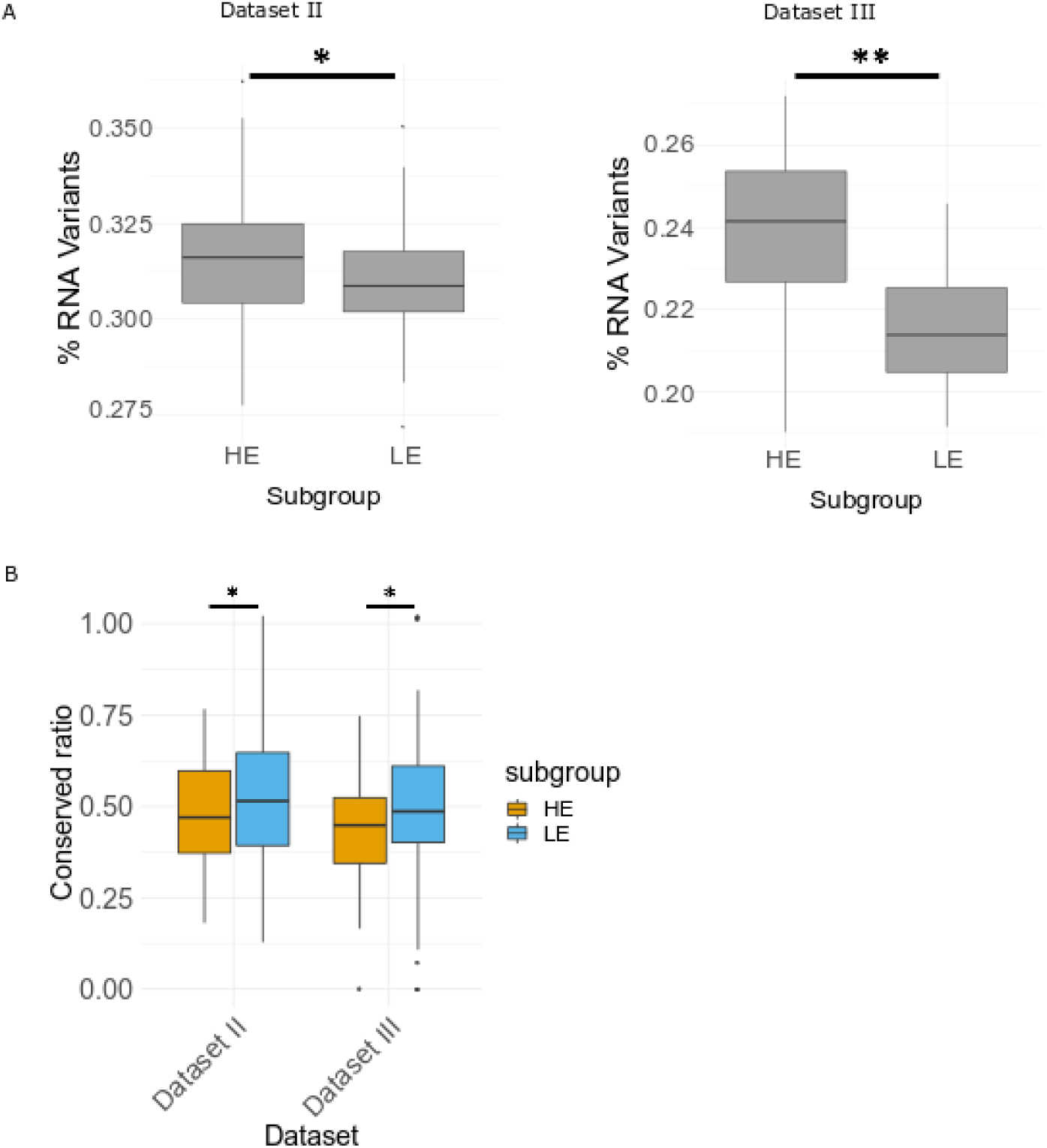
mtRNA mutation patterns display higher mutational repertoire and lower conservation score in the HE subgroup. (A) Box plot showing the comparison of the protein coding region mutational repertoire between the two subgroups, per dataset (Datasets II and III). (B) Box plot of the conserved ratios distribution of LE and HE groups, per dataset.

### Glucagon and OXPHOS genes do not consistently co-express in human pancreatic alpha cells subpopulations

As a first step to assess the generality of the distinct beta cell sub-groups to other pancreatic cell types, we took advantage of scRNA-seq data of pancreatic alpha cells from the same three datasets, considering only individuals with more than 40 alpha cells each. These criteria left us with all donors from Dataset I, seven human donors in Dataset II, including the three individuals that had sufficient numbers of beta cells; two human donors in Dataset III, excluding one individual in our above-described beta cells analysis. After applying the same approach used for beta cells analyses, although alpha cells could be divided into sub-groups according to mtDNA gene expression (Fig S11A, Fig S12, S13) they co-expressed with certain nDNA-encoded OXPHOS genes across two datasets out of three (Datasets I and II) (Fig S11B, Fig S14), yet such subgroups did not display significant expression of glucagon (GCG) between the two subgroups (Fig S11C, Fig S14). Notably, GO terms analysis revealed a weaker association with the OXPHOS and ATP metabolic processes as compared to beta cells (Table S5). In summary, although both pancreatic alpha and beta cells could be divided into subgroups according to mitochondrial gene expression, alpha cells display weaker association of nDNA-OXPHOS genes with such cellular population division, and do not correlate with their hallmark gene expression. Therefore, while considering human pancreatic islets, the association of mitochondrial gene expression with the hallmark of cellular gene expression is limited to beta cells.

### Mito-nuclear gene expression and insulin define beta cell sub-groups in new-born, but not in adult mice

We next asked whether the phenomenon of two distinct beta cell sub-groups, which are divided according to mtDNA gene expression, is conserved in evolution. The available scRNA-seq mouse datasets originate from three studies of the C57Bl6 mouse strain (termed Datasets I,II,III) (Baron et al. 2016; Xin et al. 2016a; Zeng et al. 2017), with Dataset I yielding 551 single beta cells (Baron et al. 2016), 314 single beta cells in Dataset II from 3-7 month-old mice (Xin et al. 2016a) and 387 beta cells from multiple postnatal time points in Dataset III collected from new-born mice (e.g., 84 cells collected from day 1, 87 cells from day 7, 88 cells from day 14, 68 cells from day 21, and 60 cells from day 28 postnatal) (Zeng et al. 2017). Similar to humans, mouse Dataset I was sequenced by the InDrops platform, whereas Datasets II and III were sequenced by Fluidigm C1. After quality control analysis (see Methods), 264 single beta cells remained for further analysis from mouse Dataset I, while considering ∼1600 genes genome-wide on average per cell. Filtering cells and genes in mouse Dataset II resulted in 309 single beta cells with an average of ∼5700 informative genes per cell. In mouse Dataset III (new-born mice) a total of 304 beta cells remained, including 70, 62, 69, 53 and 50 single cell for mice from day 1, day 7, day 14, day 21 and day 28, respectively, with ∼6500 informative genes on average, per cell. As in humans, the InDrops platform (Dataset I) enabled us analysing mtDNA protein-coding transcripts with >10 PolyA nucleotides (excluding Nd4l, Atp8 and Nd6; Table S6) (Ruzzenente et al. 2012). To control for high similarity (99.9%) of the mouse mtDNA sequences overlapping the genes Nd3, Nd4l, Cox2, Cox3, Atp6, Atp8 with several nuclear mitochondrial mouse pseudogenes (NUMTS) (Calabrese et al. 2012), we first limited our analysis to the seven remaining mtDNA protein-coding gene. This analysis revealed a single group of beta cells in adult mice (mouse Dataset I, II) (Fig 4A-B). In contrast, Dataset III (new-born mice) displayed two subgroups of beta cells (Fig 4C) in each of the available postnatal days, with one beta cells cluster showing higher mtDNA gene expression levels (in all tested mtDNA-encoded genes). Differential expression analysis of the orthologues nDNA-encoded OXPHOS genes in new-born mice revealed co-expression of certain structural genes, which differed among the postnatal days (Table S7). Specifically, Day 1 showed significantly high expression of certain structural and assembly genes in the LE subgroup, which were upregulated in the human HE subgroup. Day 7, and more prominently cells from days 14-28, displayed significant overexpression of certain OXPHOS structural genes in the HE group as in humans, although certain structural and assembly genes that were markers of the HE group in human were upregulated in the LE group of cells from these days. When we extended the analysis to the entire genome we found significantly higher expression of Ins2 at postnatal day 1 (p<0.001) in the LE group and higher expression of Ins1 gene (p<1×10^−5^) in the HE group. In contrast, this analysis revealed significantly higher expression of Ins2 in the HE subgroup as compared to the LE subgroup at postnatal days 14 (p<0.001), 21 (p<1×10^−5^) and 28 (p<0.05) (Fig 4D), and with significantly higher expression of Ins1 at day 14 (p<0.01) (Table S7). Finally, unlike our analysis in human beta cells, comparison of the mt-RNA mutational repertoire between the mtDNA gene expression of the beta cell clusters from the new-born mice (Dataset III) at postnatal days 14 and 28 revealed significantly higher mutational repertoire in the LE group (i.e., with the lower mtDNA gene expression) as in the tested adult mouse (p<0.005, day 14; p<1×10^−10^, day 28) (Fig S15), while day 1 and day 21 showed higher mutational repertoire in the HE subgroup as in human. Notably, similar to human, the percent of the overlapping mutations was lower than the percent of the unique mutations of the subgroups (Table S4). Nevertheless, conservation analysis of these mutations revealed that the observed ratios of each group were insignificant and inconsistent among the postnatal days (Fig 4E). It is worth noting that the results withstood a different mapping approach, namely mapping solely against the mtDNA, which enabled including all protein-coding mtDNA genes in the analysis (see methods, Fig S16, Table S4, Fig S17). In summary, these findings revealed clustering of mito-nuclear gene expression within new-born mouse beta cells, suggesting that although mt-RNA gene expression divided beta cells into subpopulation in both human and young mice, other attributes of these sub-group of cells (such as mt-RNA mutational repertoire and conservation score) diverge.

**Figure 4:**
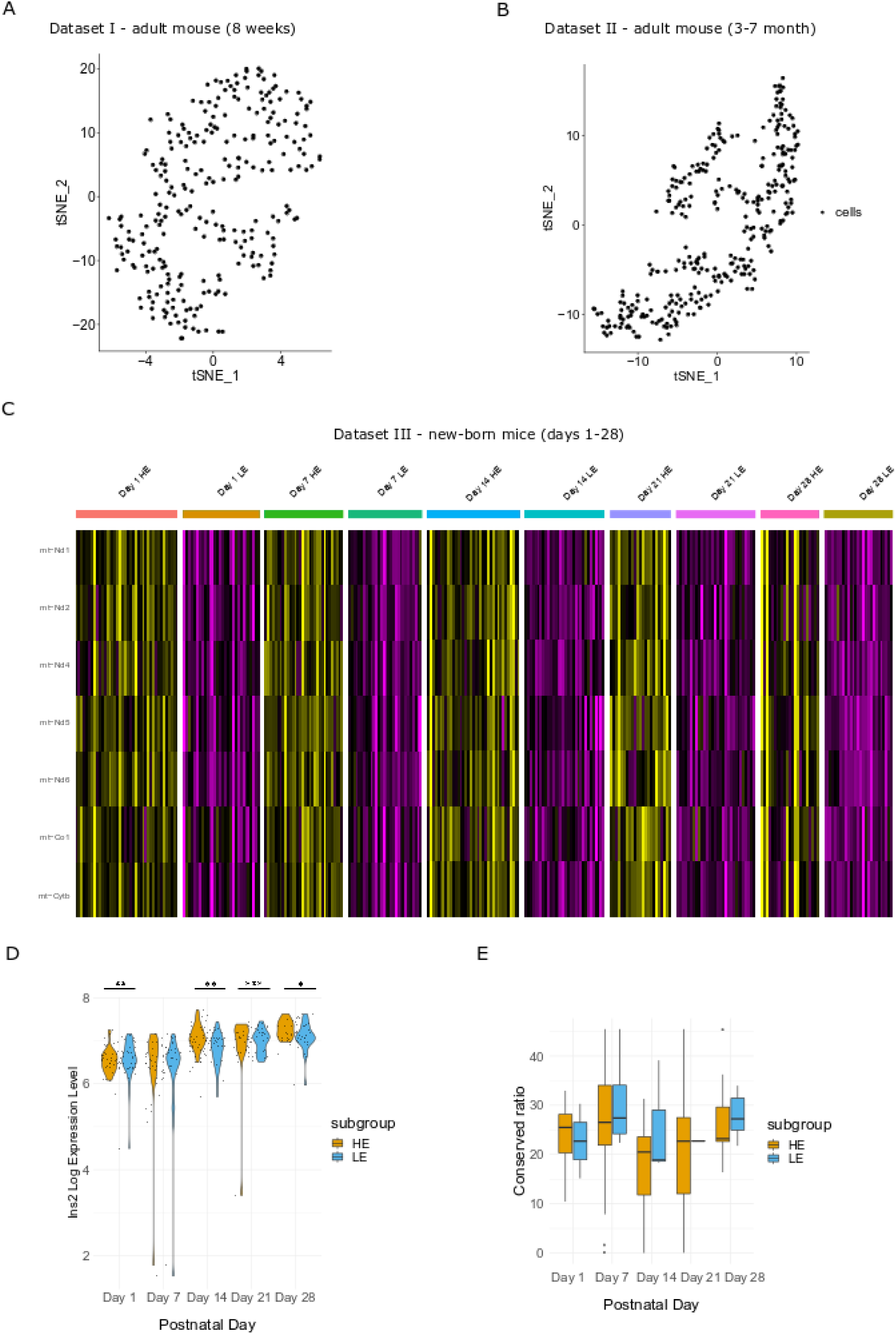
Pancreatic beta cells from newborn, but not adult mice, are divided into two clusters according to mtDNA expression. (A) tSNE profile of mtDNA gene expression pattern (protein-coding genes) in beta cells from 8 weeks old mice (mouse Dataset I). (B) tSNE of mtDNA gene expression pattern (as in A) from beta cells from 3-7 month old mice (mouse Dataset II). (C) Gene expression heatmap of mtDNA gene expression in beta cells reveal cellular sub-groups in newborn mice (1, 7, 14, 21 and 28 days postnatal – mouse Dataset III). (D) Ins2 gene expression in beta cell sub-groups in each of the tested postnatal days (mouse Dataset III). (E) Box plot of the conserved ratio of beta cells from new-born mice, per postnatal day (mouse Dataset III). Significance: * – p<0.05, ** – p<0.001, *** – p<1×10^-5^.

## Discussion

Taken together, this work revealed mitochondrial gene expression clustering in human pancreatic beta cells. Such heterogeneity was reflected by two distinct sub-populations of cells which diverged by mtDNA gene expression, nDNA OXPHOS and Insulin gene expression, and in patterns of negative selection acting on mt-RNA mutational repertoire. Since all mtDNA protein-coding genes comprise essential subunits of the OXPHOS, such differences between the sub-groups of beta cells most likely reflect previously overlooked beta cell populations divergence in terms of mitochondrial regulation, and activity. This interpretation is consistent with the positive correlation that we found with insulin gene expression. This yields a testable hypothesis – it would be of interest to test whether our observed sub-populations of beta cells correlate with beta cells activity, such as the presence of so-called insulin secreting hubs (Johnston et al. 2016) and the presence of ‘extreme’ beta cells with elevated mRNA levels of insulin versus ‘non-extreme’ beta cells that were identified in mouse (Farack et al. 2019). While human beta cells presented with a profound mitochondrial regulatory difference between two cellular sub-groups, pancreatic alpha cells did not. Specifically, although we observed an apparent sub-division into cells with different mtDNA gene expression, the expression correlation with nDNA OXPHOS genes was weaker, and the connection to the inherent function of the cell – glucagon expression, was not evident. This suggests that the mitochondrial subdivision of beta cells into subgroups is not common to all islet cell types. Furthermore, as insulin secretion has been clearly shown to rely on mitochondrial function, and alpha cells function rely more on anaerobic glycolysis (Schuit et al. 1997; Quesada et al. 2006; Mulder 2017), it is plausible that heterogeneity in mitochondrial regulation within a given cell type relies on the centrality of mitochondrial function to the tested cell type. Therefore, there is great interest in assessing mitochondrial regulatory heterogeneity in other additional cell types and tissues.

While considering mitochondrial regulatory heterogeneity in mouse beta cells, new-born mice displayed sub groups of cells which, similar to humans, diverge in their mtDNA gene expression patterns and correlated with Insulin gene expression. However, the characteristics of the subgroups in terms of nuclear gene expression changes during the development of the neonates, as days 14-28 displayed a more similar expression pattern to human as compared to days 1 and 7. This can stem from the immature metabolic phenotype of the neonatal beta cells in mice (Yoshihara et al. 2016). In contrast, while considering mitochondrial gene expression, the adult mice (8 weeks and 3-7 month) displayed a more homogenous population of beta cells. The observed differences between human and adult mouse beta cells might stem from the islets architecture (Cabrera et al. 2006) and the difference in longevity of the beta cells (Cnop et al. 2010). Thus, it will be of interest to explore whether the beta cell mitochondrial sub-groups in humans also appear in children, and if they do, whether they correlate with expression of nDNA-encoded mitochondrial genes, as well as with the expression of Insulin.

The identification of the human beta cells subgroups in both healthy and type 2 diabetes individuals support the fundamental importance of such subpopulations of cells for life. As the sample size of humans tested is relatively small, future increase in sample size is required to draw any conclusion about the implications of our observations to disease conditions. With this in mind, it would be of interest to eventually study beta cells mitochondrial subdivision of gene expression in type I diabetes patients, and see whether any of the identified subgroups is more prone to dysfunction in patients.

The positive correlation of mtDNA gene expression in the identified beta cells subgroups with insulin gene expression lends a first clue for the physiological importance of these subgroups. Specifically, our findings suggest that human beta cells diverge into functionally different groups already at the gene regulatory level, and not only physiologically (Johnston et al. 2016). Nevertheless, it still remains to be found whether the nature of the subgroups and their composition will change upon exposure to mitochondria-related environmental conditions. Previous reports demonstrated the contribution of oxidative stress and reactive oxygen species (ROS) to insulin secretion in response to changing glucose levels, and in embryogenesis (Leloup et al. 2009; Hoarau et al. 2014). Notably, the latter is also correlated with the expression of the transcription factor Jun-D (Laurent et al. 2008), which has recently been shown not only to regulate gene expression in the nucleus but also be imported into human mitochondria and bind the mtDNA (Blumberg et al. 2014). These findings tempt us to ask whether manipulation of JunD, or other factors that are candidates to modulate the bi-genomic regulation of gene expression, will alter the composition of the two beta cells groups and their response to certain environmental conditions. These future experiments will enable better understanding of the role of beta cells mitochondrial heterogeneity in health and in disease conditions.

## Online Methods

### Available samples for analysis

scRNA-seq data from mouse and human pancreatic beta cells were obtained from five studies. Datasets were downloaded from the following sites;

Human and mouse Dataset I: https://www.ncbi.nlm.nih.gov/geo/query/acc.cgi?acc=GSE84133

Human Dataset II: https://www.ncbi.nlm.nih.gov/geo/query/acc.cgi?acc=GSE81608

Human Dataset III: https://www.ncbi.nlm.nih.gov/geo/query/acc.cgi?acc=GSE86473

Mouse Dataset II: https://www.ncbi.nlm.nih.gov/geo/query/acc.cgi?acc=GSE77980

Mouse Dataset III: https://www.ncbi.nlm.nih.gov/geo/query/acc.cgi?acc=GSE86479

### Processing of scRNAseq data

For Dataset I of human and mouse, the bioinformatics pipeline of the data processing was carried out as previously reported (Baron et al. 2016).

For Datasets II and III of human and mouse sequenced reads were trimmed using Trim Galore (version 0.4.5; https://www.bioinformatics.babraham.ac.uk/projects/trim_galore/) while employing default parameters, in addition to the following parameters: [--clip_R1 n] and [--three_prime_clip_R1 n] (n – representing 5% of the read length) to avoid low quality bases and potential adapter contamination. Trimmed reads were mapped against the reference human genome (GRCh38 for human cells and GRCm38 for mouse cells) using STAR (version 2.5.3) (Dobin et al. 2013). Mapping of the sequencing reads in the human datasets was performed using default parameters, in addition to the [—outFilterMultimapNmax 1] parameter, to achieve unique mapping, as previously performed (Cohen et al. 2016) to avoid contamination of expressed mitochondrial pseudogenes – mtDNA fragments that were transferred to the nucleus during the course of evolution (NUMTs, see dedicated section below) (Mishmar et al. 2004). Non-unique mapping was also performed for comparison and assessment of such potential contamination. As mouse NUMTs were longer and more similar to the active mtDNA, sequencing reads from six mtDNA genes were erroneously filtered out while applying the unique mapping protocol. To overcome such problem,, sequencing reads from the mouse datasets were mapped solely against the mtDNA genome using *bwa* with *aln* parameter (BWA-backtrack algorithm) (Li and Durbin 2009); this enabled subsequent analysis for all mtDNA encoded-genes. Expression levels of all genes were counted using HTSeq-count v0.11.2 (Anders et al. 2015), using default parameters and employing the [-f bam] parameters. For quality control filtering, gene count values as defined by HTSeq-count were concatenated into a resulting gene expression matrix for each library, which then was loaded into R for subsequent computational analysis. Seurat objects were created using the function “CreateSeuratObject” (Butler et al. 2018). Additionally, for quality control filtering of Dataset I (inDrops platform), cells and genes were filtered as previously reported (Baron et al. 2016) In brief, for further analyses we used cells having at least 3,000 detected transcripts, with a maximum of 20% ribosomal genes. For all datasets cell doublets were excluded (i.e., cells that were assigned to a given cell type – beta or alpha cell, yet express a mixture of cell type-specific markers – both Glucagon and Insulin those) (Fig S1, S2). Cells with either more than 15% mtDNA reads, or less than MEAN-2.5*SD % mitochondrial reads or with zero inflated mtDNA reads (in at least mtDNA-encoded gene) were removed. These measures were taken since overrepresentation of mtDNA genes expression could either associate with stress, and with cell death (Ilicic et al. 2016).

### Cluster identification using Seurat

To identify cluster of pancreatic beta cells which share patterns of mitochondrial gene expression, Seurat pipeline was utilized (Butler et al. 2018). The data matrices were imported and processed with Seurat R package version 3.0.2. To account for the possibility that individual cell complexity leads to cluster separation and subsequent reduction in the number of total read counts per cell, we used the “vars.to.regress” parameter in scaling function of Seurat. PCA was performed for each separate individual (for both human and mouse experiments) using the mtDNA-protein coding mRNA genes. Although the mtDNA codes for 37 genes, of which 13 encode essential protein-subunits of the oxidative phosphorylation (OXPHOS) system, 2 rRNA genes (12S, 16S) and 22 tRNA genes, the RNA-seq libraries of all datasets enabled analysis of only longer transcripts, while excluding transcripts with short 3’ poly-A (i.e. <10A) in the inDrops platform, which selected for PolyA+ transcripts (Dataset I) (Slomovic et al. 2005). This limited our analysis to the 13 mtDNA-encoded protein coding genes for the Fluidigm C1 platform and to 9 of the 13 mtDNA-encoded OXPHOS subunits (excluding ND5, ND6, ND4L, ATP8 which have a short polyA tail) in the inDrops platform (Table S6). Although the mtDNA is transcribed in strand-specific polycistrons, it is not obvious that mtDNA transcripts will express in the same levels mainly due to post-transcription processing; therefore, multidimensional clustering was performed. Using the first two principle components as input, density clustering was performed per individual to identify groupings in the data and t-distributed statistical neighbour embedding (tSNE) to visualize. A range of values (0.1-1) were examined to assess differences in mitochondrial gene expression. To gain statistical power, the cells of all individuals were clustered and the percent of cells that were consistent with their group identity was calculated and these cells were used for subsequent analyses. Using further Seurat functionality applications, marker genes for each respective cluster were identified and used for subsequent analysis. The specific markers for each cluster identified by Seurat were determined using the “FindAllMarkers” function, using only highly expressed genes (non-zero genes above 0.25 of cells).

### Statistical analyses

Statistical analysis for categorical groups comparisons was performed by unpaired Wilcoxon test with Jack knife re-sampling test that was performed to ensure uneven groups comparisons are correct with 1000 test repeats. Mutation repertoire and conserved ratio differences were tested using ANOVA test. Differential expression of genes was tested using “negbinom” test for Dataset I which identifies differentially expressed genes between every two groups by performing a likelihood ratio test of negative binomial generalized linear models and the “bimod” test for Datasets II and III which developed for measurements from the Fluidigm platform (McDavid et al. 2013). Differential expression of genes was corrected with FDR correction for multiple comparison.

### Mitochondrial sequence extraction

Bam files were indexed using default parameters of Samtools v1.3.1 (index command). To create multiple sequence alignment, we generated pileup files using the Samtools mpileup command (default parameters). In addition, we used the -r MT parameter in order to determine read counts per cell, per-base; to facilitate the usage of this parameter for each studied sample we used a custom-made Python script for each sequenced sample. For each given mtDNA position, with sufficient read coverage that passed our quality control filters (see below), the base frequency was calculated by dividing the number of reads which display a certain base by the total coverage per position.

### Variant quality control and filtering

We counted base changes (RNA mutations), only in mtDNA positions covered by at least 400 sequencing reads. A mutation was considered trustworthy only if it was covered by at least two sequencing reads from each direction (e.g., forward and reverse) and if the identified mutation was not in the end of the sequencing read. Secondly, high quality variants per sample, per position were determined if the total coverage of the position was >400 with the mutation represented by >1% coverage in a given nucleotide position. To avoid low coverage errors, only cells with at least 1000 covered mtDNA positions in the protein coding region were included in the variant analyses. Due to low coverage in the non-coding mtDNA region (D-loop), only mutations in the mtDNA coding region were used for subsequent comparison of mutational repertoire between cells. While considering the mtDNA mutational repertoire, bootstrap analysis (with 1000 repeats) was performed by resampling 1000 high quality positions per iteration, per cell. Mutations percentage per subgroup was calculated by summing the variants per subgroup and dividing by 1000 X #of cells per iteration.

### Identification of personalized sub-group mutations

To identify mutations that are more prevalent in one group of cells as compared to the other per individual, or per tested condition, frequencies of mutations and heteroplasmy percentage (mean plus SD) were determined in each of the cell groups (Table S4). Additionally, candidate inherited mutations were identified if they were shared between alpha and beta cells isolated from the same individual. For each individual the percent of candidate inherited mutations was determined by dividing the number of such by the total number of mutations, per cell subgroup, per individual.

### Assessing the functional potential of mutations in mtDNA transcripts

To assess whether RNA mutations occurred randomly throughout the mtDNA, or were subjected to constraints, the conservation score averages of all the detected position was compared to random distribution. To this end, 100-way phastCons (Siepel et al. 2005; Pollard et al. 2010) score per human and mouse mtDNA position was downloaded from the UCSC website (http://genome.ucsc.edu/), and the average score of all RNA heterogenic positions was calculated for each sample. The scores of random distribution were calculated by sample-specific bootstrapping. For each sample, the original number of detected heterogenic positions was resampled ten thousand times, and the average score of all the resampled positions in each iteration was calculated. Next, the expected random value was calculated by averaging the score of all iterations. The ratio between the observed score average, and the expected random average, was calculated to compare between the distributions of the two cellular sub-populations, per subject. Sample specific p-values were calculated based on the bootstrap scores, as the fraction of iterations that had either lower or higher score averages than the observed average (when the observed-expected ratio was lower or higher than 1, respectively).

### Mitochondrial nDNA pseudogenes (NUMTs) likely did not impact expression differences

It has been known for some time, that nDNA harbors a repertoire of mtDNA sequence fragments (NUMTs) that were transferred from the mitochondria during the course of evolution. NUMTs potentially pose an obstacle to mtDNA gene expression assessment, as a subset of RNA reads might originate from NUMTs rather than from the active mtDNA. As a first step to control for such a scenario, we performed both unique and non-unique mapping in the human datasets. Similarly, in *mus musculus* there is a large NUMT covering a substantial part of the mtDNA (∼4.5 kb), with high sequence similarity to the corresponding mitochondrial reference genome (99.9% across this sequence) (Calabrese et al. 2012). As leaving this part out will result in data loss for 6 mtDNA genes in mouse, we used *bwa* mapping only for the mtDNA genome sequence. To test the levels of potential NUMTS in the unique mapping data, the percent of NUMT reads (+/−SD) were calculated per cell, per base (Table S8). In addition, whole genome differential expression analysis further filtered out pseudogenes to avoid noise.

## Acknowledgements

This study was funded by the Israel Science Foundation grant number 372/17 and by the US Army life sciences division grant number LS67993 awarded to DM. The authors thank Dr. Tal Shay (BGU) for critical discussion during early stages of manuscript preparation.

## Supplementary information

### Tables S1-S8

Table S1 – Details of individuals.

Table S2 – Comparisons of cell type and health status.

Table S3 – Marker gene expression of beta cells subgroups.

Table S4 – mtRNA mutations per human subject/mouse postnatal day.

Table S5 – Marker gene expression of alpha cells subgroups.

Table S6 – Human and mouse polyA tail of mtDNA transcripts.

Table S7 – Marker gene expression of new-born mouse beta cells subgroups.

Table S8 – percentage of NUMT reads (+/− SD).

### Figures S1-S17

Figure S1 – Violin distribution plots of expression of selected marker genes in pancreatic alpha and beta cells, in Dataset I.

Figure S2 – Violin distribution plots of expression of selected marker genes in pancreatic alpha and beta cells, in Datasets II and III.

Figure S3 – Beta cells display higher mtDNA gene expression regardless of diabetes status in Dataset I.

Figure S4 – Beta cells have higher mtDNA gene expression regardless of diabetes status (non-unique mapping) in Datasets II and III.

Figure S5 – Beta cells have higher mtDNA gene expression regardless of diabetes status (unique mapping) in Datasets II and III.

Figure S6 – Beta cells from Datasets II and III (unique mapping) diverged according to mtDNA gene expression into high (HE) and low (LE) subgroups.

Figure S7 – Beta cells from Datasets II and III (non-unique mapping) diverged according to mtDNA gene expression into high (HE) and low (LE) subgroups.

Figure S8 – Human mtDNA gene expression analysis revealed two distinct beta cell clusters with either high or low mtDNA gene expression (assigned as HE and LE, respectively).

Figure S9 – Plots showing lack of association between the expression of nuclear antigens CD9 and ST8SIA1 in the LE and HE cellular subgroups.

Figure S10 – Human mtRNA mutation patterns display a tendency towards higher mutational repertoire and lower conservation score (see Online Methods) in the HE subgroup in each of the tested six individuals.

Figure S11 – Pancreatic alpha cells are divided into two sub-groups according to mtDNA gene expression.

Figure S12 – Pancreatic alpha cells are divided into two sub-groups according to mtDNA gene expression (non-unique mapping).

Figure S13 – Pancreatic alpha cells are divided into two sub-groups according to mtDNA gene expression (unique mapping).

Figure S14 – Heatmap reveals that pancreatic alpha cells are divided into two sub-groups according to mtDNA gene expression.

Figure S15 – mtRNA mutational pattern analysis in mouse datasets- unique mapping.

Figure S16 – Adult mice (8 weeks and 3-7 month) did not show subgroups of beta cells, while new-born mice displayed two subgroups of beta cells according to mtDNA gene expression.

Figure S17 – mtRNA mutational pattern analysis- bwa mapping (mapping solely against the mtDNA genome) of mouse Dataset III.

